# The influence of incompatibilities and heterosis on hybrid population genetics

**DOI:** 10.64898/2026.08.04.742766

**Authors:** Julio A. Ayala-López, Stephan Peischl, Claudia Bank

## Abstract

A long-standing question in evolutionary biology is: Under what circumstances can speciation occur despite hybridisation or because of hybridisation? Some models of hybrid incompatibilities predict that speciation can occur even in the presence of gene flow and strong selection against hybrids, an outcome also influenced by genetic contributions from parental species and genetic architecture. On the other hand, empirical work has shown that heterosis can counteract the effect of incompatibilities, hindering the speciation process. Theoretical models that simultaneously consider the positive and negative impacts of hybridisation on fitness remain scarce, raising questions about the effects of hybrid incompatibilities in the presence of heterosis. To address this question, we study how (Bateson)-Dobzhansky-Muller incompatibilities (BDMIs) interact with overdominant mutations in a two-locus population genetics model of an isolated hybrid population. We find that the strength of overdominance relative to incompatibilities determines the long-term genetic composition of the hybrid population. We show that high recombination exposes incompatibilities to selection and reduces the frequency of derived alleles in the hybrid population, limiting the strength of BDMIs that can be maintained by the balance with overdominance. We also show how neutral variation is affected by the strength of selection and the recombination rate between incompatible loci, generating patterns that include an increase in local variation resembling associative overdominance, or a reduction resembling background selection. Such variation of neutral variation, particularly at intermediate distances from BDMI loci, can generate peaks or troughs of diversity that are explained by the recombination rate between BDMI loci, and initial proportions of admixture between parental populations. Our work demonstrates how the genomic conflict caused by the interplay of overdominance and hybrid incompatibilities, recombination, and parental contributions, shape the genome of an isolated hybrid population.

## Introduction

Hybridisation, the interbreeding of previously isolated populations, plays a role in several evolutionary processes, such as adaptive radiations (Seehausen, 2004), adaptive introgression (Rieseberg & Wendel, 1993), and, centrally, speciation (Abbott et al., 2013). However, even when reproductive isolation evolves, it often remains incomplete and is permeable, as shown with growing evidence of gene flow between closely related species (Mallet, 2005). Understanding the conditions that allow populations to hybridise –and their genomic signatures and consequences– requires a theory that acknowledges the interaction between mechanisms that prevent or facilitate gene flow.

A common observation in experimental crosses is that F1 or F2 crosses exhibit lower fitness than their parental populations. The (Bateson-)Dobzhansky-Muller model is the standard model for explaining this form of intrinsic postzygotic isolation (Bateson, 1909; Dobzhansky, 1936; Muller, 1942). This model proposes that substitutions accumulate at different loci as populations diverge in isolation. An incompatibility occurs when substitutions from different populations co-occur in a hybrid genome, resulting in detrimental epistatic interactions between alleles that have not encountered each other during their establishment. The resulting hybrid incompatibilities, known as (Bateson)-Dobzhansky-Muller incompatibilities (BDMIs), have been widely studied in the context of reproductive isolation, hybrid zones, and gene flow (Bank et al., 2012; Cutter, 2012; Gavrilets, 1997; Orr, 1995; Orr & Turelli, 2001). One prediction is that, in parapatry, BDMI loci must be adaptive in each population to maintain a barrier (Bank et al., 2012).

Contrastingly, hybridisation can sometimes be beneficial, and hybrid offspring can have greater fitness than their parents, a phenomenon known as heterosis (Shull, 1948). Theoretical and empirical work suggests that heterosis can influence patterns of divergence and introgression by transiently masking deleterious mutations or facilitating the spread of adaptive alleles across populations (Dagilis et al., 2019; Schneemann & Welch, 2023). Such fitness advantages may affect reproductive isolation depending on genetic architecture, ecological context and the tight balance between heterosis and incompatibility, at least in early-generation hybrids (Thompson & Schluter, 2022). The study of heterosis is extensive in the context of crop-improvement (Reviewed in Labroo et al., 2021), and most theoretical work focuses on transient heterosis in early generation hybrids; however, the consequences of heterosis on an evolutionary timescale remain poorly understood (but see Schneemann & Welch, 2025).

Although there have been considerable advances over the last few decades in understanding the genetic basis of hybrid incompatibilities and heterosis (Schneemann & Welch, 2025; Schumer et al., 2015), these phenomena are mainly studied separately. Crucially, hybrid incompatibilities and heterosis are expected to have opposing signatures of selection on the genomic landscape. Strong selection against BDMIs should locally reduce genetic diversity, whereas heterosis in the form of overdominance can increase linked diversity (Ohta & Kimura, 1970; Pamilo & Pálsson, 1998; Zhao & Charlesworth, 2016). As distinct processes can generate heterogeneous genomic landscapes of differentiation between populations, it remains a challenge to identify how these processes generate different signatures of selection (Reviewed in Ravinet et al., 2017). Here, we examine a minimal model of a BDMI that incorporates heterosis via overdominance. We investigate how the interaction of hybrid incompatibilities and heterosis can sustain and shape an isolated hybrid population. We quantify the role of recombination in modulating genomic conflict at BDMI loci and characterise the expected patterns of linked neutral variation. This work provides a theoretical account for the maintenance of hybrid allele combinations without the need for ongoing gene flow.

## Methods

### Model description

We model a scenario in which a single admixture event between two locally adapted populations generates a new, isolated hybrid swarm. Following secondary contact, the hybrid population evolves without any subsequent gene flow from the parental populations. The evolution of this hybrid population is governed by selection via overdominant heterosis and negative epistasis in the form of BDMIs, and by recombination. Specifically, we analyse a two-locus model of a diploid, panmictic population with loci *A* and *B*. Ancestral alleles are denoted by lowercase letters and derived alleles by uppercase letters. We consider two isolated parental populations: one fixed for haplotype *Ab*, and the other fixed for haplotype *aB*. Following secondary contact, these populations contribute proportions 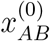 and 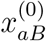 to a new isolated hybrid population (Fig. 1A). Selection acts on diploid zygotes, with fitness as specified in Table 1.

**Table 1:** Two-locus fitness matrix with overdominance determined by parameters *α, β*, and epistasis determined by parameters *γ*_*i*_.

|  | aa | aA | AA |
| --- | --- | --- | --- |
| bb | 0 | $\alpha$ | 0 |
| bB | $\beta$ | $\alpha + \beta - \gamma_1$ | $\beta - \gamma_2$ |
| BB | 0 | $\alpha - \gamma_3$ | $-\gamma_4$ |

**Figure 1:**
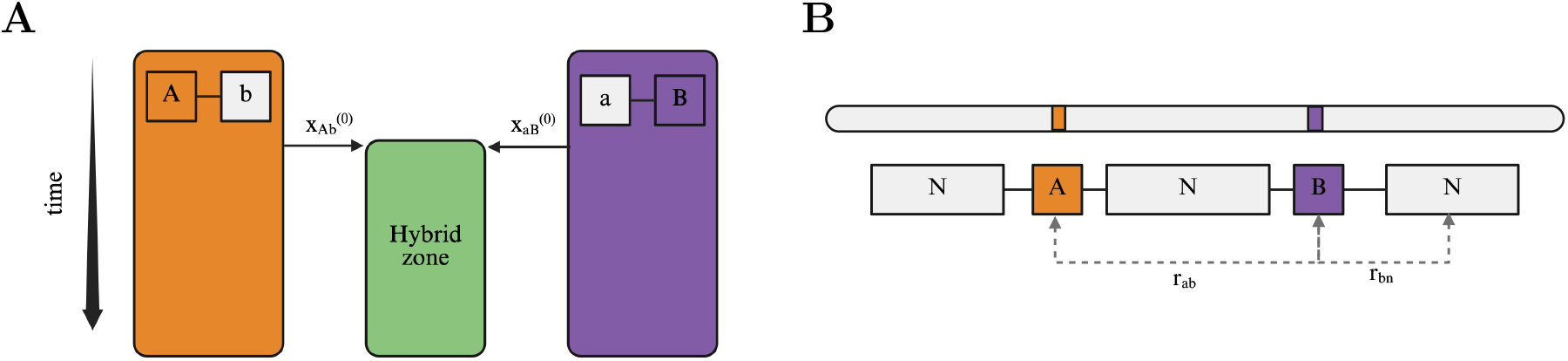
Representation of demographic scenario and genetic architecture **A**. Initially, populations *Ab* (orange) and *aB* (purple) exist in isolation. Then, a hybrid swarm is formed by a proportion 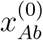 of individuals from population *Ab*, and a proportion 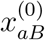 from population *aB*, followed by isolation, resulting in a single hybrid population with no subsequent gene flow. **B** Schematic representation of the genetic architecture, where a BDMI between *A* and *B* with recombination rate *r*_*ab*_, is surrounded by an arbitrarily positioned neutral locus *N. B* recombines with *N* at rate *r*_*bn*_.

Heterosis is modelled as overdominance (i.e. heterozygote advantage), where heterozygote individuals for *A/a* have a fitness advantage *α* over homozygotes, and *B/b* have a fitness advantage *β*. We assume that overdominance is symmetric, i.e. it has the same effect at both loci (*α* = *β*).

We model hybrid incompatibilities as negative epistatic interactions between derived alleles, i.e. as BDMIs, distinguishing between homozygous and heterozygous interactions following the model of Turelli and Orr (2000). The negative interaction is quantified by para meter Γ = *{γ*_1_, *γ*_2_, *γ*_3_, *γ*_4_*}*, where *γ*_*i*_ is the fitness cost imposed on genotypes *AaBb, AABb, AaBB, AABB*, respectively. In the main text, we focus on a co-dominant model of BDMIs with fitness costs proportional to the number of conflicts between derived alleles: Γ = *{γ/*2, *γ, γ*, 2*γ}*, reducing epistasis to a single parameter *γ*. In the supplementary material, we include results for recessive BDMIs with no fitness costs for double heterozygotes (*γ*_1_ = 0).

We model evolution of the population by tracking the frequencies of the four haplotypes *ab, Ab, aB, AB* as a vector ***x*** = *{x*_1_, *x*_2_, *x*_3_, *x*_4_*}* : *x*_*i*_ ≥ 0 and 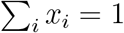. By assuming continuous-time dynamics, the haplotype dynamics of the system are described by the set of equations:

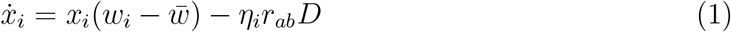

where 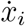 is the change in haplotype frequency over time, *w*_*i*_ is the marginal fitness, 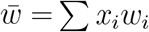 is the Malthusian mean fitness, *η*_1_ = *η*_4_ = −*η*_3_ = −*η*_2_ = 1, *r*_*ab*_ is the recombination rate between *A* and *B*, and *D* = *x*_1_*x*_4_ − *x*_2_*x*_3_ is a measure of linkage disequilibrium. Expressions for marginal and mean fitness are provided in the supplementary material

(Eq. S1, S2). Eq. 1 can be rewritten in terms of allele frequencies:

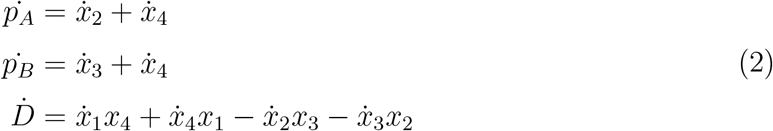

To study the evolutionary consequences of the interaction between overdominance and BDMIs in the hybrid population, we quantified the parameter range in which derived alleles coexist over long timescales. We focus on scenarios in which polymorphisms at both loci exist at equilibrium (*p*_*A*_ *>* 0 and *p*_*B*_ *>* 0). We first study limiting cases of recombination: loose linkage, where loci are assumed to segregate independently; and no recombination between incompatible loci (*r*_*ab*_ = 0). We then analyse the model for intermediate recombination values. Moreover, we study the dynamics at a neutral locus neighbouring the BDMI in a three-locus model (See supplementary material for model description). Analytical and numerical analyses were performed in Wolfram Mathematica 13.3. Numerical simulations for the linked selection model were done in Python 3.11.8. All scripts and notebooks are available at https://github.com/banklab/DMI-heterosis and will be archived on Zenodo upon publication.

## Results

### Maintenance of BDMI alleles in the hybrid population

#### High recombination

If recombination is sufficiently large relative to selection, we can assume that loci are independent and in linkage equilibrium (LE). In the absence of BDMIs, polymorphism at both *A* and *B* is favoured due to overdominance. As an incompatibility is added, such polymorphism can persist only if the strength of BDMIs is low relative to overdominance, as negative epistasis selects against recombinant *AB* haplotypes. A fully polymorphic equilibrium maintaining both *A* and *B* alleles in the population is possible, with allele frequencies given by

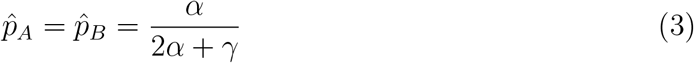

Due to symmetry in overdominance (*α* = *β*), 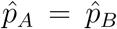 when a fully polymorphic equilibrium is reached. This equilibrium is asymptotically stable if 0 ≤ *γ <* 2*α*, that is, full polymorphism can be maintained if the strength of BDMIs is less than twice the strength of overdominance. When the strength of BDMIs is higher (*γ* ≥ 2), the BDMI is resolved (i.e. one of the BDMI alleles is lost), resulting in one of two single-locus polymorphic equilibria: 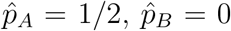, and 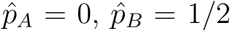, maintained by overdominance, and reached depending on initial admixture proportions, as illustrated in Fig. S1B.

At the limit of quasi-linkage equilibrium (QLE), a first-order approximation of allele frequencies and linkage disequilibrium is given by:

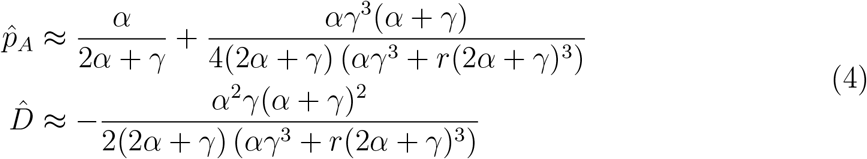

Eq. 4 shows the contribution of strong recombination to the increase in frequency of derived alleles, as *r* is always positive and the contribution is in the order of 1*/r*. LD is always negative, as *x*_*Ab*_ and *x*_*aB*_ haplotypes are overrepresented in the population, and approaches 0 in the LE limit as *r* increases.

### Low recombination

The previous section demonstrates that recombination exposes incompatible alleles to selection, limiting the coexistence of incompatible alleles in the hybrid population in scenarios where overdominance is sufficiently strong to offset the cost imposed by BDMIs. If recombination between BDMI loci is reduced, incompatibilities appear less often in the same background, leading to negative LD. To consider the case of low recombination, we first begin with the limiting case of complete linkage between *A* and *B* (*r* = 0), reducing the system to a four-allele haploid model. Here, one-locus polymorphisms exist when *A* or *B* go extinct, as the BDMI is resolved. Additionally, there is one fully polymorphic equilibrium described by:

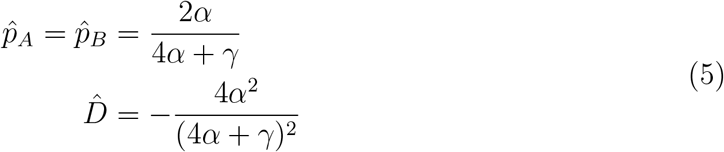

This equilibrium is asymptotically stable if 0 ≤ *γ <* 4*α*, allowing incompatible alleles to be present in the population when BDMIs are up to 4 times as strong as the effect of overdominance. Notably, 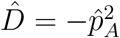 ; haplotypes *x*_*AB*_ are removed from the population due to their detrimental effect –and not generated by lack of recombination–, maximizing repulsion LD. For the limiting cases of LE and no recombination, 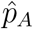 is reduced from a maximum of 1*/*2 when *γ* = 0, converging to 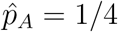 as *γ* → 2*α* for LE, and *γ* → 4*α* for no recombination (Fig. 2C, lines).

**Figure 2:**
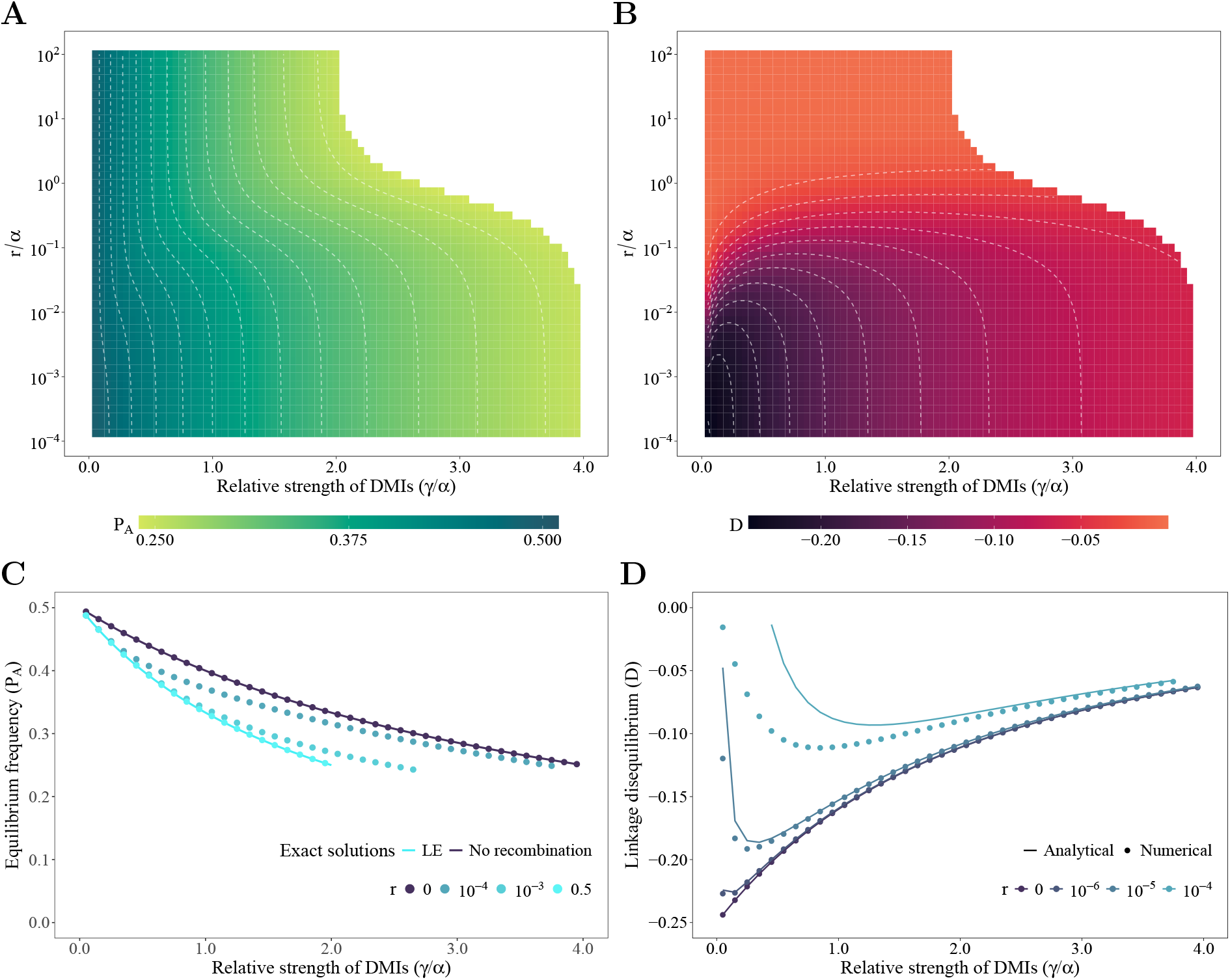
Maintenance of fully polymorphic hybrid populations (filled area, **A, B**) depends on the interaction of the strength of co-dominant BDMIs relative to overdominance (*γ/α*), and the recombination rate, *r*. **A** Equilibrium frequency of derived alleles (*p*_*A*_ = *p*_*B*_) with relative strength of BDMIs to overdominance (*γ/α*), x-axis, and relative recombination rate to overdominance (*r/α*), y-axis. **B** Linkage disequilibrium, *D*, as a function of *γ/α*, and *r/α*. Dotted white lines are used as a visual guide for similar values. **C** Equilibrium frequency of derived alleles, *p*_*A*_, *p*_*B*_, as a function of (*γ/α*) for different values of *r*, represented for numerical solutions (dots), and exact solutions for limiting cases (lines). **D** Linkage disequilibrium, *D*, as a function of (*γ/α*) for different values of *r*, represented for numerical solutions (dots), and analytical approximations from Eq. 6 (lines). All results shown for values of *α* = 0.01.

A first-order approximation for low recombination (0 *< r* ≪ 1) based on (Eq. 5) for the equilibrium frequency and linkage disequilibrium is:

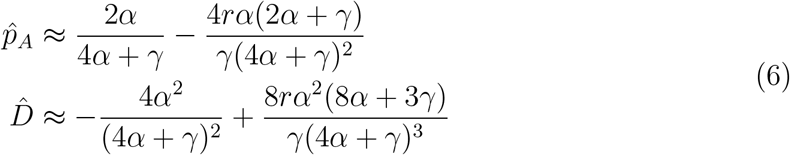

Here, the equilibrium frequency, 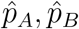, depends not only on the strength of BDMIs and overdominance (*γ/α*) as shown before, but is also reduced by recombination. Consistently, recombination contributes to the erosion of linkage disequilibrium, as the correction term in *r* is positive, decreasing the magnitude of LD, as illustrated in Fig. 2D.

### General model behaviour

For intermediate recombination, our model exhibits behaviour between the limits shown in the previous sections. To find the regions of the parameter space where full polymorphism can be maintained, as well as allele frequencies and linkage disequilibrium, we numerically solve the dynamical system for a grid of parameter combinations, given by *r* and the strength of BDMIs relative to the strength of overdominance *γ/α*. We find that the limits of relative strength of BDMIS when polymorphism is lost decrease non-linearly with increasing recombination (Fig. 2A). The frequencies of derived alleles at equilibrium decrease with increasing strength of BDMIs. Notably, LD does not change monotonically as the change in frequency does with increasing *γ/α*. When both recombination and the strength of BDMIs are minimal, LD is strongest, as recombination is weak compared to the strength of overdominance, favouring *Ab* and *aB* haplotypes (Fig. S5B). This non-monotonic pattern of LD is particularly observed at low recombination rates from Eq. 6, exemplified in Fig. 2B,D.

## Linked variation at neutral sites

Selection also affects genetic diversity at linked neutral loci. Neutral variation can either decrease due to background selection (BGS) caused by the BDMI, or increase due to associative overdominance (AOD) caused by overdominant heterosis. To quantify this effect, we study a three-locus model on a neutral locus *N*, positioned at varying distances relative to the two selected loci *A* and *B*. We assume that the neutral allele *N* is initially fixed in one of the parental populations and absent from the other. In other words, when *N* is present in a *B* genetic background, the hybrid population is initially composed of haplotypes *aBN* and *Abn* with frequencies 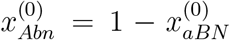, where *N* can either be positioned after locus within or around selected loci *A* and *B* (see 1). This gives a neutral expectation of heterozygosity determined by the initial admixture proportions. Linked selection distorts neutral heterozygosity away from this expectation, calculated at equilibrium at the neutral site (*Ĥ*_*N*_). We show the conditions in which, selection at *A* and *B* increase or decrease neutral diversity based on the position of *N*, and on parental contributions 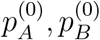.

### Neutral heterozygosity at flanking regions of epistatic loci

If overdominance is the only selective force (i.e., with no BDMI), a neutral allele *N* hitchhikes, generating AOD at neighbouring sites. Linked neutral diversity is then maximised close to the selected loci and decays following a sigmoid pattern. Thus, the equilibrium heterozygosity ranges from *Ĥ*_*A*_ = *Ĥ*_*B*_ = 0.5 when completely linked, to the initial heterozygosity of the neutral locus, 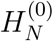, as the distance from the nearest overdominant locus increases (Fig. 3A). For neutral loci located between *A* and *B*, this pattern is only visible under sufficiently strong recombination (*r*_*ab*_ = 0.1 in Fig. 3A). If *A* and *B* are located more closely to each other, this pattern disappears, as overdominance is stronger than recombination (*r*_*ab*_ = 0.01; *r*_*ab*_ = 10^−^6 in Fig. 3A, centre panel).

**Figure 3:**
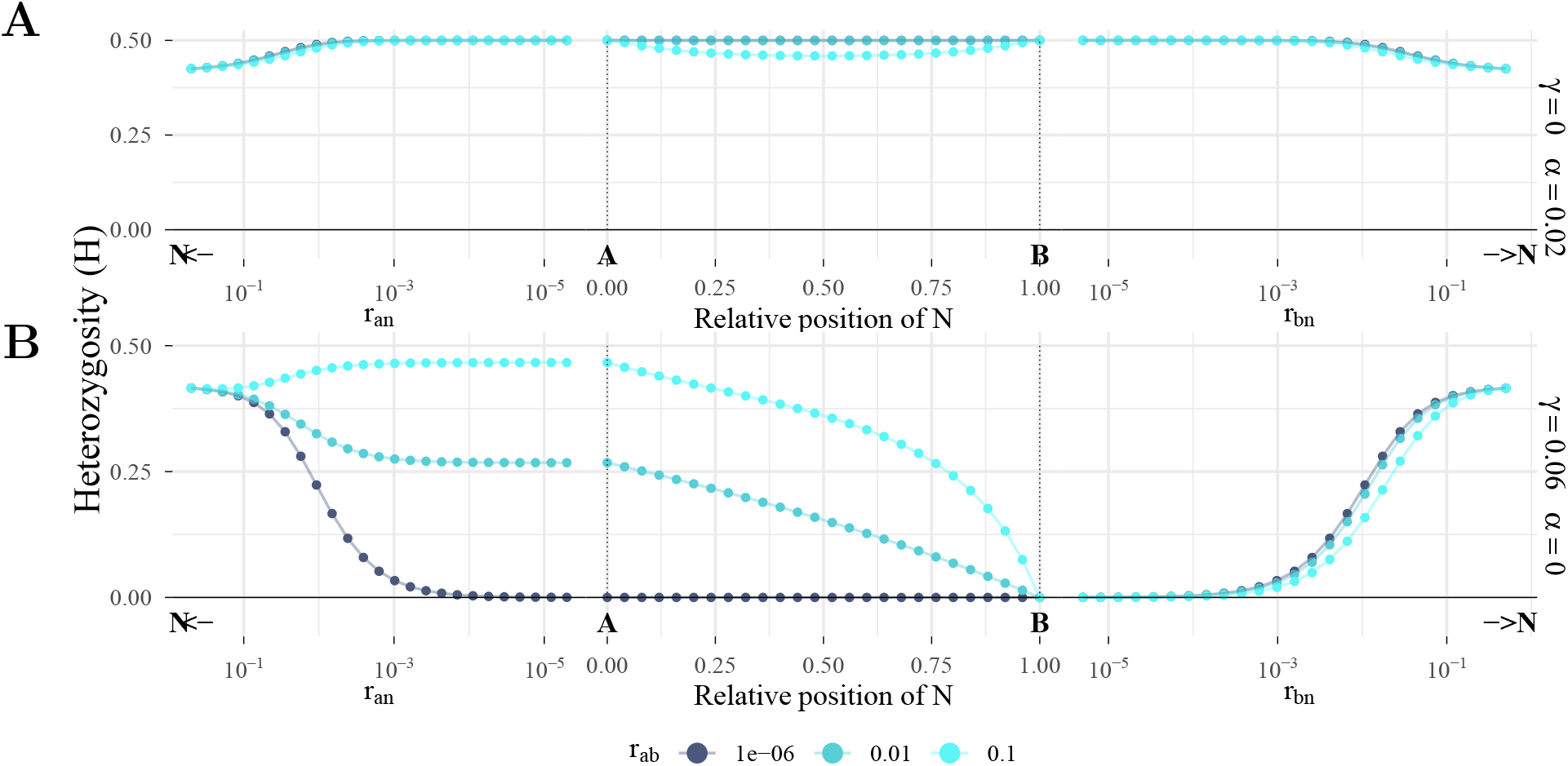
Patterns of heterozygosity at a neutral locus linked to overdominant loci or codominant BDMIs vary with distance from selected loci and the strength of selection. **A** Top row: Neutral heterozygosity across a sequence linked to overdominant loci (*α* = 0.02). **B** Bottom row: Neutral heterozygosity across a sequence linked to a co-dominant BDMI (*γ* = 0.06). Initial allele frequencies: 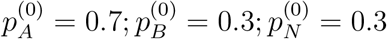.

When there is a single co-dominant BDMI and no overdominance (*α* = *β* = 0), the minor parental allele always goes extinct (e.g., *B* in the example of Fig. 3B), and the BDMI is always resolved. During this process, the major allele *A* decreases in frequency due to epistatic selection and becomes neutral when *B* is lost. This change in frequency is reflected as a change in *Ĥ*_*A,B*_ relative to 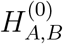 (Fig. S4A,B). The equilibrium heterozygosity at locus *A, Ĥ*_*A*_, increases with recombination distance *r*_*ab*_ (coloured large dots, center panels in Fig. 3), as the BDMI is resolved faster. Neutral loci flanking the BDMI monotonically change in heterozygosity between the equilibrium heterozygosity at the selected loci and their initial heterozygosity, 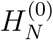, to which it converges with increasing recombination. Here, the width of the diversity trough is affected by the recombination distance between BDMI loci (Fig. 3B, right panel), as well as the strength of the BDMI (Fig. S6).

We next consider a scenario where a BDMI and overdominant heterosis occur simultaneously, with *Ĥ*_*A,B*_ given by the equilibrium frequencies described in the general model behaviour section. The signature on neutral heterozygosity varies broadly depending on the recombination rate of the neutral locus, the strength of selection on BDMIs relative to overdominance (*γ/α*), and the initial frequency of the neutral allele 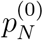. When *N* is positioned on the left side of the sequence, linked more strongly to *A*, sites at intermediate recombination distance increase in heterozygosity relative to 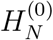 and *Ĥ*_*A*_ (Fig. 4A, left panels) if *B* does not go extinct due to a strong BDMI. Conversely, when *N* is positioned on the right side of the sequence, linked to *B, Ĥ*_*N*_ at intermediate recombination distance can slightly decrease relative to *Ĥ*_*B*_ when the strength of BDMIs is intermediate (Fig 4A, purple line), or when recombination is intermediate and BDMIs are strong (Fig 4B, red line). Interestingly, neutral variation within the BDMI can also be higher than *Ĥ*_*A,B*_ at low recombination distance when BDMIs are strong (Fig. 4C, red line), generating AOD. Variation within the BDMI can be reduced closer to the minor parent *B* with higher recombination rates (Fig. 4A, purple line). In this scenario, when *N* is more closely linked to *A* than *B*, it can increase *Ĥ*_*N*_ . However, the increase does not necessarily occur where *N* is positioned halfway between *A* and *B*, as the pattern depends on the asymmetry in initial frequencies 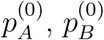.

**Figure 4:**
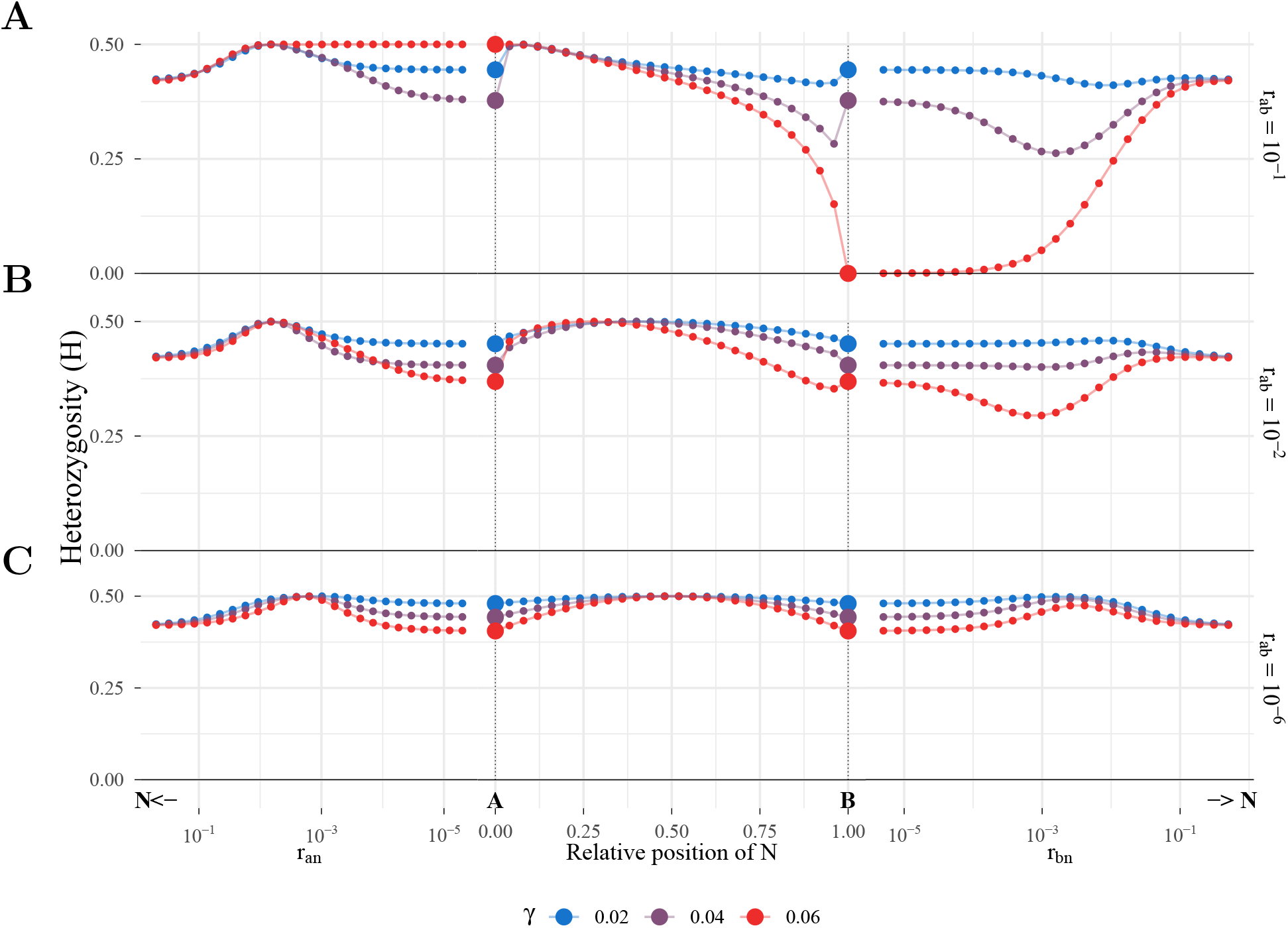
Patterns of heterozygosity at a neutral locus linked to a co-dominant BDMI when incompatible loci *A* and *B* are both overdominant (*α* = *β* = 0.02), with varying recombination *r*_*ab*_ between incompatible loci (high to low, from up to bottom rows). **A** Top row: High recombination (*r*_*ab*_ = 10^−^1) leads to the extinction of *B* if the strength of the BDMI is high (large red dots), or lead to different patterns of heterozygosity with distance. **B** Middle row: Intermediate recombination (*r*_*ab*_ = 10^−2^) leads to a non-monotonic change in heterozygosity with varying strength of BDMIs. **C** Low recombination *r*_*ab*_ = 10^−6^) can lead to an increase in neutral heterozygosity relative to equilibrium or unlinked expectations. Initial allele frequencies for all scenarios: 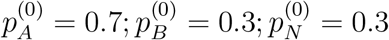 ;

## Discussion

In this work, we describe the conditions under which a BDMI is maintained in an isolated hybrid population in the presence of overdominant heterosis. Upon hybridisation, incompatible alleles in a BDMI become exposed to selection, typically leading to the resolution of the incompatibility in the absence of gene flow, regardless of the strength of selection. Our minimal two-locus model shows that overdominance at the same loci can lead to stable, long-term maintenance of BDMIs in an isolated hybrid population. The maintenance of polymorphism depends strongly on the recombination rate between the interacting loci. Here, we derive analytical expressions for the expected frequency and linkage disequilibrium between BDMI loci in the limits of no recombination and LE, as well as approximations for QLE and low recombination. Moreover, we describe the effect of arbitrary recombination between BDMI loci numerically, and describe the signature of a BDMI with overdominance at linked neutral loci, where we find a combination of patterns of AOD and hitchhiking.

When recombination is sufficiently weak, BDMI alleles can be maintained if they are up to twice as strong as the total overdominance at both loci (4*α* if *α* = *β*). The absence of recombination allows incompatible alleles to coexist in different genetic backgrounds, favouring polymorphism at a single locus. Note that if *A* and *B* are completely linked, this polymorphism only exists if the *ab* haplotype is initially present in the hybrid population, possible by the ancestral haplotype segregating in the population or through back mutation, though we do not consider the role of mutation in this work. If only *Ab* and *aB* haplotypes are initially present in the population and recombination between *A* and *B* is absent, an unstable equilibrium of *p*_*A*_ = *p*_*B*_ = 0.5 exists, as double heterozygotes *Ab/aB* have an advantage over the homozygotes *Ab/Ab* or *aB/aB*.

Because the maintenance of hybrid allele combinations relies on restricted recombination, our findings align with theoretical expectations that strong negative epistasis favours modifiers that suppress recombination (Lenormand & Otto, 2000). In our work, reduced recombination favours the coexistence of BDMI alleles, as overdominance confers an advantage greater than the cost of BDMIs, provided there are enough ancestral haplotypes to maintain this balance. In fact, in the limit of no recombination and maximal strength of BDMIs allowing complete polymorphism (*γ* → 4*α*), an equilibrium is reached with haplotype frequencies 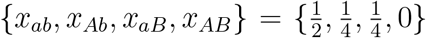 (Fig. S5A). In natural populations, recombination suppression frequently occurs within chromosomal inversions, which can capture and shield BDMI loci from breaking down, even under gene flow (Feder & Nosil, 2009; Ortiz-Barrientos et al., 2016). Empirically, the capture of BDMIs within inversions is a well-documented driver of speciation and hybrid maintenance, as classically observed in Drosophila (Noor et al., 2001; Poikela et al., 2024). Overdominant selection, as modelled here, provides a robust mechanism to permanently stabilise incompatibilities within such regions. In this work, we treat the recombination rate between BDMI loci as a fixed parameter. Future theoretical work modelling the evolution of recombination explicitly through a modifier would allow us to test whether and how selection drives the evolution of recombination between a BDMI in the presence of overdominance.

When recombination is present, the generation of the recombinant haplotype *AB* exposes the population to the fitness cost of the incompatibility. The analytical solutions for linkage and quasi-linkage equilibrium show that the maximum relative strength of BDMIs to overdominance drops from 4*α* down to 2*α* as recombination increases. This shift in the threshold for maintenance of incompatible alleles shows that the genetic architecture of an incompatibility is a primary determinant of its lifespan upon secondary contact: loosely linked BDMIs act as stronger, but shorter-lived, barriers to gene flow, whereas tightly linked BDMIs act as weaker, but longer-lived, ones (Fig. S4C,D). High recombination continually exposes the BDMI to selection, reducing derived allele frequencies of incompatible loci (Fig. 2C) and driving the incompatibility toward its resolution as the relative strength of the BDMI increases (*γ* → 2*α*; Fig. 2A). Tightly linked BDMIs, by contrast, are exposed to selection only rarely, so the barrier they present is weaker, but the polymorphism can be maintained up to twice the relative strength of the BDMI (*γ* → 4*α*). As loosely linked loci are rapidly purged of their incompatibilities via recombination, reproductive isolation due to BDMIs is likely to be caused by tightly linked epistatic loci (Barton & Bengtsson, 1986; Butlin, 2005), shielded in regions of reduced recombination (Boman et al., 2025; Schumer et al., 2018), even in the presence of over-dominant heterosis (Eq. 6, Fig. 2). Reduced recombination prevents introgression by maintaining hybrid alleles at low frequencies in separate backgrounds, even when BDMIs are strong, as reflected by negative LD (Fig. 2C, D). The maintenance of incompatible alleles, even when selection against them is strong, is contingent on overdominance acting directly at the BDMI loci. We make this simplifying assumption to isolate overdominance as a direct cause for the maintenance of a BDMI. If overdominance occurs at different loci, the BDMI could be resolved, with a speed depending on the recombination distance between BDMI and heterotic loci. We do not consider this more general scenario here, leaving it as a natural extension of the present model.

Because the maintenance or resolution of a BDMI is coupled with its recombination rate, the conflict between overdominance and epistasis leaves a distinct, heterogeneous footprint in the surrounding genome. Notably, our analysis of linked neutral variation demonstrates that neutral genetic diversity at loci flanking a BDMI is variable and strongly modulated by recombination-selection dynamics and initial admixture proportions. The epistatic interaction in the BDMI generates patterns of heterozygosity that deviate from the signature of hitchhiking expected at a single selected locus (Fig. 3B; cf. Kim and Stephan, 2002). A BDMI alone generates a pattern comparable to recurrent hitchhiking (Wiehe & Stephan, 1993) or a soft sweep (Hermisson & Pennings, 2005), as selection acts on pre-existing variation to purge maladaptive epistatic combinations. This process reduces local variation and generates a broad trough of diversity, a direct consequence of the high initial frequency of the maladaptive allele. To estimate the strength of selection that generates such a hitchhiking pattern, we fit a model of recurrent hitchhiking (Wiehe & Stephan, 1993) to our numerical solutions of a BDMI where the incompatibility gets resolved by the extinction of the minor parental allele (Fig. S8A). We show that the estimated selection coefficient depends, naturally, on the strength of selection against the BDMI, *γ*, and the recombination rate, *r*_*ab*_, between BDMI loci (Fig. S8B). By replicating hitchhiking patterns through epistatic selection, we highlight a caveat for genomic scans of selection that rely solely on patterns of nucleotide diversity. Because a trough of diversity can arise from multiple processes including demographic history, background selection (Charlesworth & Jensen, 2021), and epistasis, the inferred strength of selection should be interpreted with caution, as different processes can generate similar patterns of diversity.

Heterozygote advantage, on the other hand, can generate patterns of associative over-dominance (Ohta & Kimura, 1970; Pamilo & Pálsson, 1998). When a BDMI interacts with overdominance, we find that depending on the distance to the barrier loci and the initial frequency of the parental haplotypes, distinct peaks or troughs of neutral diversity can be observed. The stable maintenance of the BDMI by overdominance elevates linked neutral diversity neighbouring the major allele at an intermediate distance (Fig. 4, left panels). Contrastingly, a trough of diversity can be observed neighbouring the minor parental allele if recombination is high between the BDMI (Fig. 4, top and middle right panels). The opposite is observed when recombination is low (Fig. 4, bottom right panel), where an increase in neutral variation is observed at intermediate recombination distance. Similarly, an increase in neutral variation at intermediate distances is observed on the left side of the major parental allele at intermediate distances, more apparent with increasing strength of BDMIs (Fig. 4, left panels). This increase in neutral variation generates a pattern of AOD originating not only from overdominance, but also from selection against deleterious allele combinations originating from the BDMI. This pattern of AOD originating from negative selection has previously been studied in regions of low recombination (Becher et al., 2020; Gilbert et al., 2020; Marion & Noor, 2023). In our work, this pattern of AOD can be observed at intermediate recombination distance when the proportion of initial admixture between parental populations is asymmetric (i.e., *p*_*A*0_ ≠ *p*_*B*0_), andis consistent in recessive BDMIs when *r*_*ab*_ is low (Fig. S3). When admixture proportions on secondary contact are equal, the pattern of elevated linked neutral diversity disappears, and AOD is observed only between the incompatibility (Fig. S7).

The sensitivity to initial admixture proportions highlights how parental contributions shape the genomic landscape of a hybrid population (reviewed in Moran et al., 2021). Previous theoretical work has found that the probability of hybrid speciation is maximised when admixture proportions are close to equal (Blanckaert & Bank, 2018), and BDMI detection is sensitive to the deviation from equal parental contributions (Li et al., 2022). Empirically, introgression from the minor parental species has been found in regions of high recombination, as observed in swordtail fish (Schumer et al., 2018). Our results provide additional context for these results: high recombination between BDMI loci exposes them to selection, reducing diversity neighbouring the minor allele and creating diversity troughs(Fig. S4C). Similarly, the diversity neighbouring the major parental allele remains high, as recombination acts faster than selection on the BDMI (Fig. S4B).

In this work, we highlight the effects of uneven parental contributions to the hybrid population and different patterns of neutral variation surrounding a BDMI. Neutral heterozygosity is initially equal to the initial heterozygosity of the *A* and *B* loci, an assumption we make to facilitate interpretation of our results. Future work could relax several of our simplifying assumptions to better approximate the fate and genomic signatures of hybrid populations. First, heterosis in hybrid populations can arise from multiple sources beyond overdominance, and different models such as pseudo-overdominance arising from the masking of recessive deleterious mutations fixed in parentals could be studied. Incorporating this source of heterosis would allow us to differentiate the patterns left on linked neutral diversity from the ones generated by a BDMI. Second, our model assumes infinite population sizes and monomorphic parental populations; relaxing this assumption by, for example, modelling unequal *N*_*e*_ between parental populations would allow to clarify how prior demography shapes linked neutral diversity neighbouring a BDMI. Furthermore, extending beyond a single pulse of admixture to include continuous migration, spatial structure, or post-admixture bottlenecks, would help us to assess how the signatures we describe here are affected in more complex hybrid populations. Finally, developing models that incorporate ongoing gene flow is a natural next step toward testing these predictions against empirical data from well-described systems, such as swordtail fish (Schumer et al., 2018) or house mice (Turner & Harr, 2014), would help determine their detectability. Comparing the patterns described here to those observed in natural populations would help determine the detectability of these signatures under more realistic demographic scenarios.

Our results contribute to the broader question of how genetic architecture and dominance interactions shape patterns of diversity and divergence. Classic analyses by Pamilo and Pálsson (1998) demonstrated that dominance variation and heterozygote advantage can generate substantial population-level heterozygosity even in the presence of weak selection. While we restrict our analysis to a single BDMI, our model adds to the study of complex hybrid incompatibilities by coupling a BDMI with overdominant heterosis. Overall, our model highlights that the genomic footprint of reproductive isolation cannot be interpreted independently of the local recombination rate and the selective forces acting within and between BDMI loci. Ultimately, unravelling the evolutionary fate of admixed populations will require moving beyond simple models of epistatic conflict to explicitly account for complex recombination-selection interactions.

## Data and code availability

All scripts and notebooks used for analysis and plotting are available at https://github.com/banklab/DMI-heterosis and will be archived on Zenodo upon publication.

## Author contributions

JAL and CB conceived the idea. JAL wrote the model, code, and the first draft of the manuscript. All authors contributed to the methodology, analysis, reviewing and editing of the manuscript.

## Supplementary material

### Model

The dynamical equations from the main text (Eq. 1) have marginal fitness for all haplotypes:

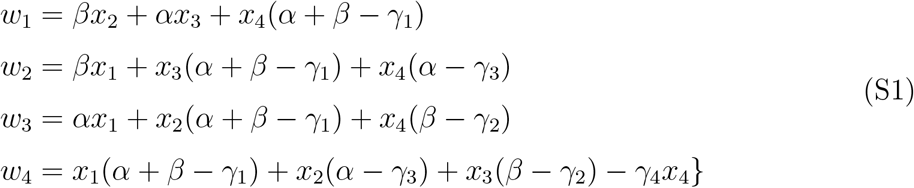

and mean fitness 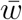 given by

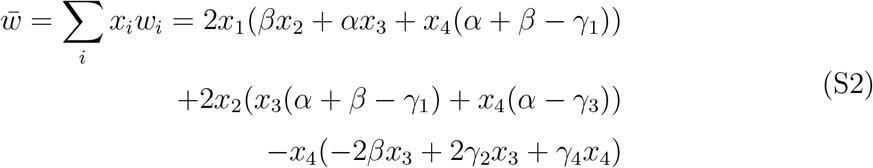

### Variation at a third neutral locus model

An extension of the two-locus model adds a third, neutral, biallelic locus *N* that can be positioned after or between *A* and *B* (1B). The addition of this third locus introduces a second recombination parameter *r*_*bn*_ when *N* is positioned after *B*. Dynamics for haplotypes *abn, Abn, aBn, ABn, abN, AbN, aBN, ABN* with corresponding frequencies 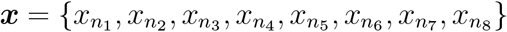 is described by the system of equations:

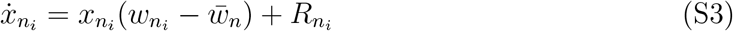

where the *n* denotes the dynamical system with a neutral locus, and 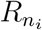describes the change in frequency of 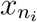due to recombination:

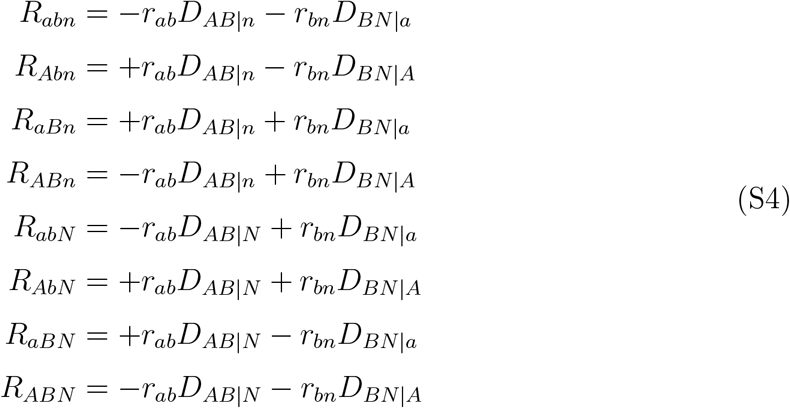

With expressions for linkage disequilibrium conditional on background

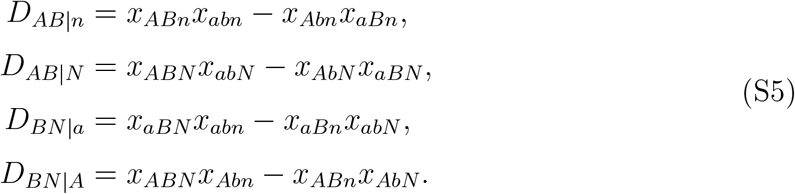

As selection does not act on the neutral locus directly, selection acts equally in all *N* haplotypes regardless of their allelic value (e.g. *w*_*abn*_ = *w*_*abN*_). Moreover, if selection is weak, the system is equivalent to a model in discrete time with multiplicative fitness (Nagylaki, 1992). The frequency dynamics of all haplotypes, analogous to Eq. S3, can be expressed by:

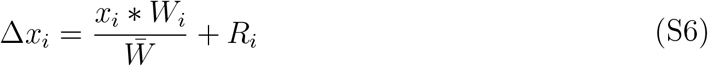

where 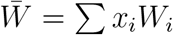 is the mean fitness and *W*_*i*_ is the marginal multiplicative fitness based on Table S1,

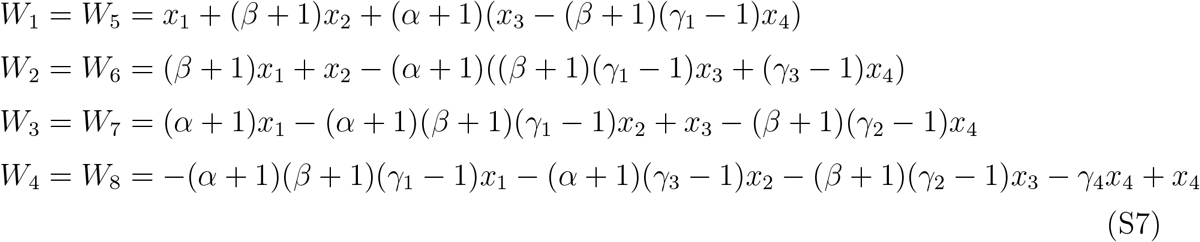

**Table S1:** Multiplicative fitness matrix for two loci.

|  | aa | aA | AA |
| --- | --- | --- | --- |
| bb | 1 | $1 + \alpha$ | 1 |
| bB | $1 + \beta$ | $(1 + \alpha)(1 + \beta)(1 - \gamma_1)$ | $(1 + \beta)(1 - \gamma_2)$ |
| BB | 1 | $(1 + \alpha)(1 - \gamma_3)$ | $1 - \gamma_4$ |

### Phase portraits

**Figure S1:**
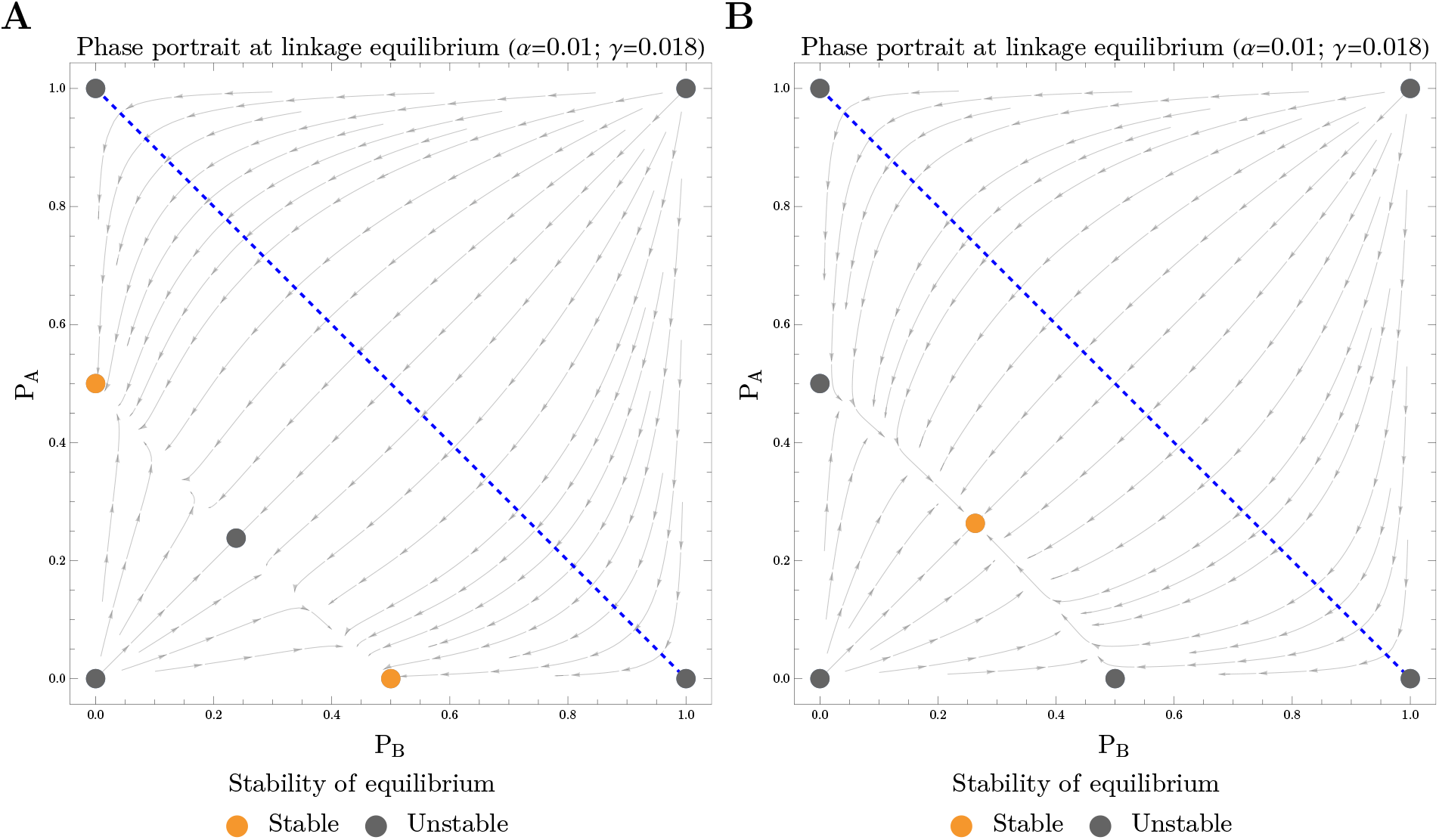
Vector fields for the change in allele frequency when the system is in linkage equilibrium with varying strength of co-dominant BDMIs. **A** There is a single asymptotically stable solution when the strength of overdominance is high relative to BDMIs (*α* = 0.01; *γ* = 0.018). **B** If BDMIs are too strong (i.e. *γ >* 2*α*), the internal fully polymorphic equilibrium becomes unstable. Two single-locus polymorphisms at *p*_*A*_ = 1*/*2, *p*_*B*_ = 0 and *p*_*A*_ = 0, *p*_*B*_ = 1*/*2 become stable, reflecting a shift in stability. Dashed blue lines indicate the range of initial frequencies for both populations.

### Recessive BDMIs

**Figure S2:**
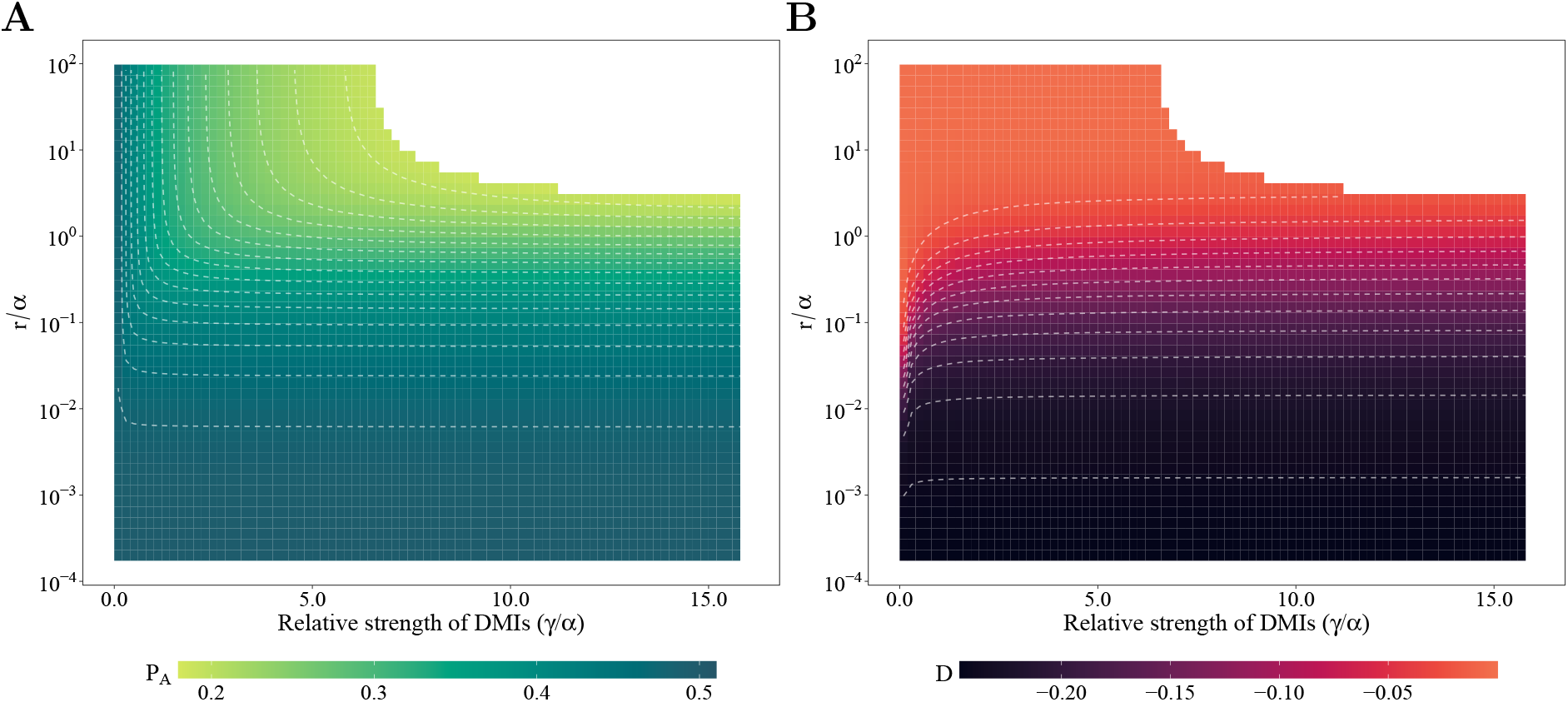
Maintenance of fully polymorphic hybrid populations (filled area) with over-dominance and recessive BDMIs. Parameters: 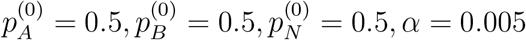

**Figure S3:**
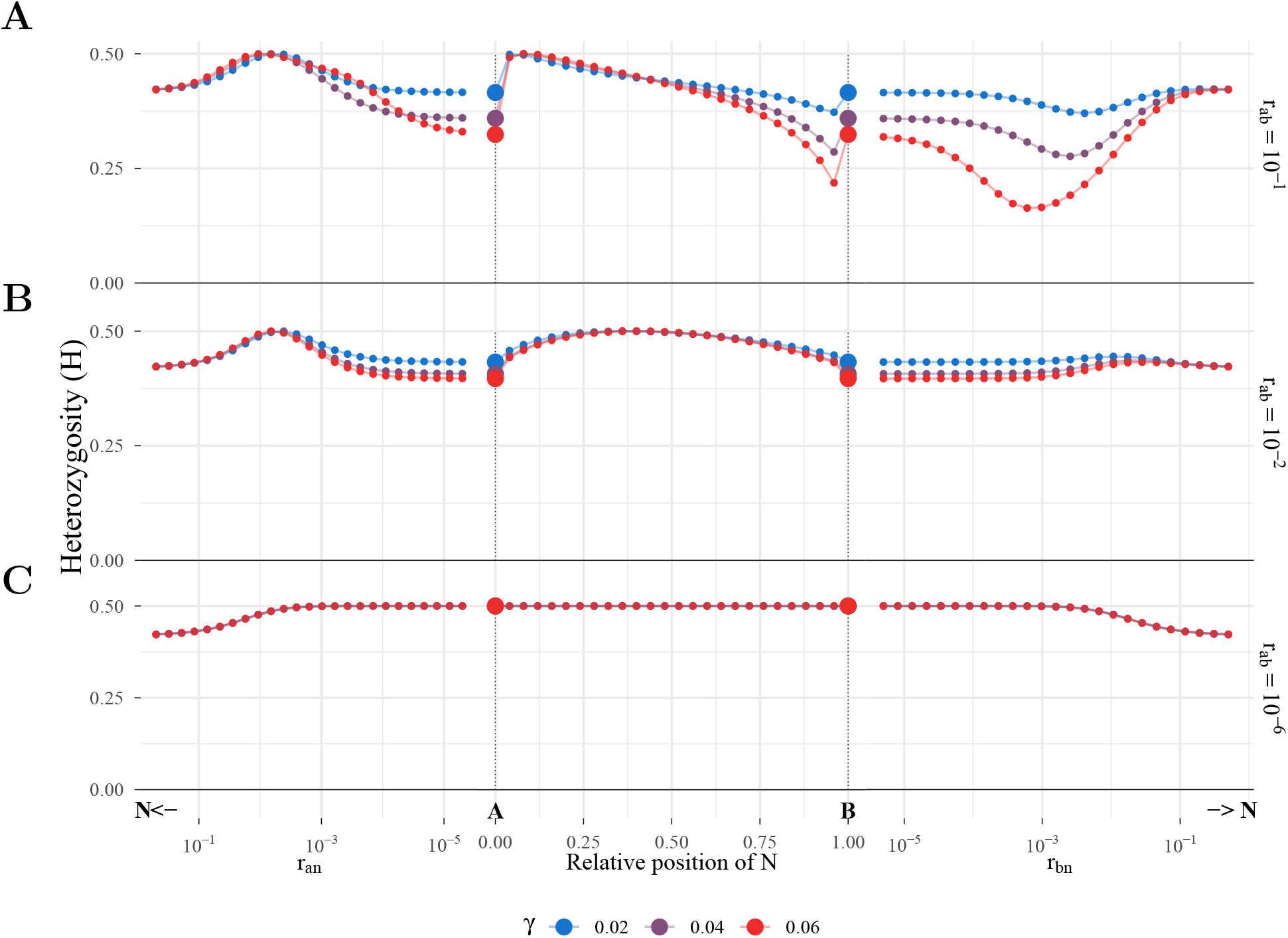
Maintenance of fully polymorphic hybrid populations (filled area) with over-dominance and recessive BDMIs. Parameters: 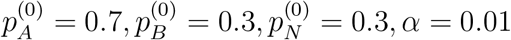

### Temporal dynamics

**Figure S4:**
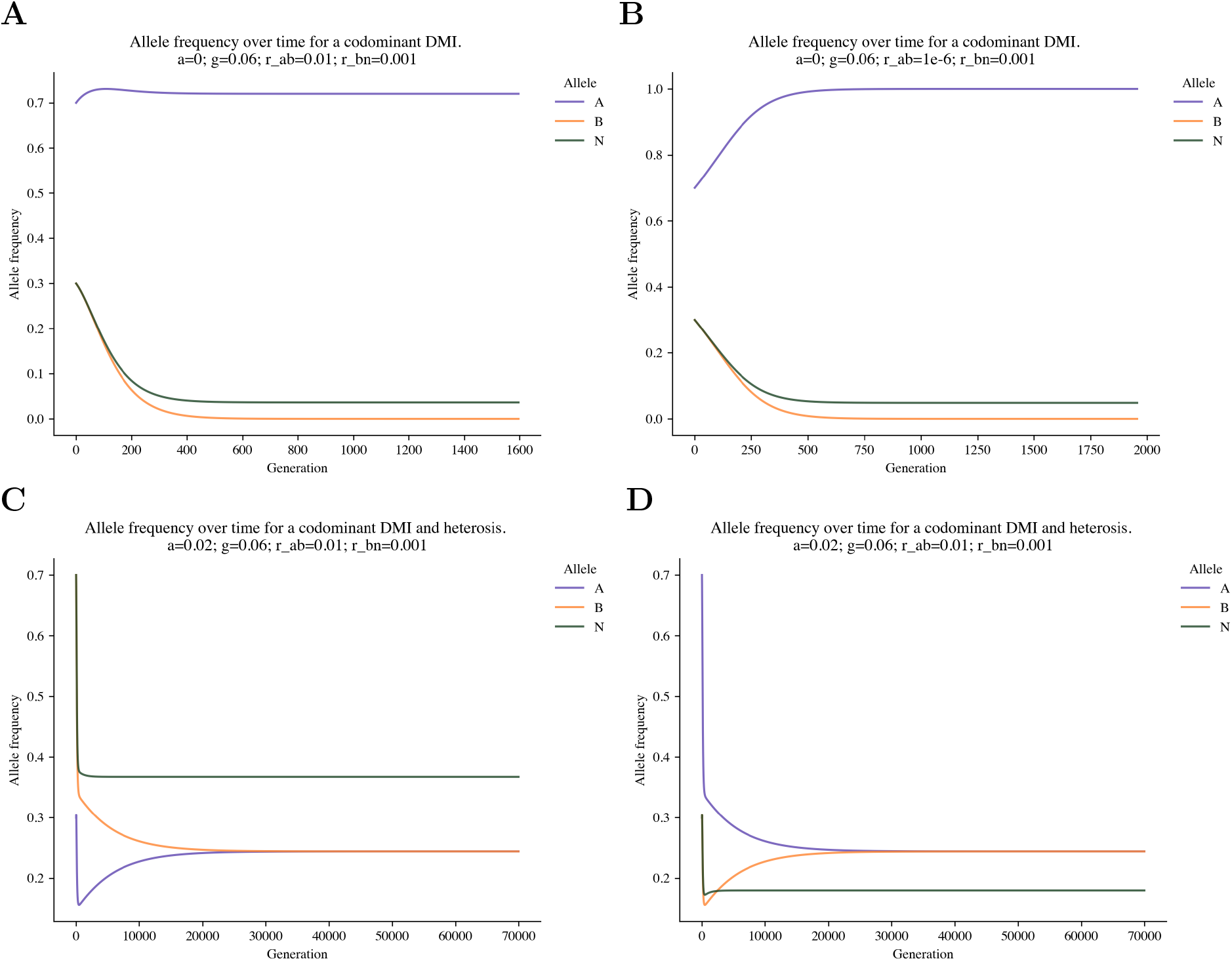
Allele frequency trajectories over time until equilibrium is reached on a co-dominant BDMI, and on neutral allele *N*. **A**,**B** Recombination distance affects the frequency of major parental allele (A) at equilibrium before the minor parental allele (B) goes extinct. **A** *r*_*ab*_ = 0.01.**B** *r*_*ab*_ = 10^−6^**C** Equilibrium of incompatible alleles when overdominance is present reaches the same equilibrium due to symmetry. The frequency of neutral *N* remains high due to recombination while *B* is decreasing in frequency, resulting in higher neutral than BDMI loci heterozygosity. 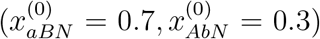 **D** If *B* and *N* start at low frequencies 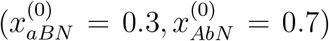, *N* further decreases in frequency before recombination acts, reflected as lower heterozygosity.

**Figure S5:**
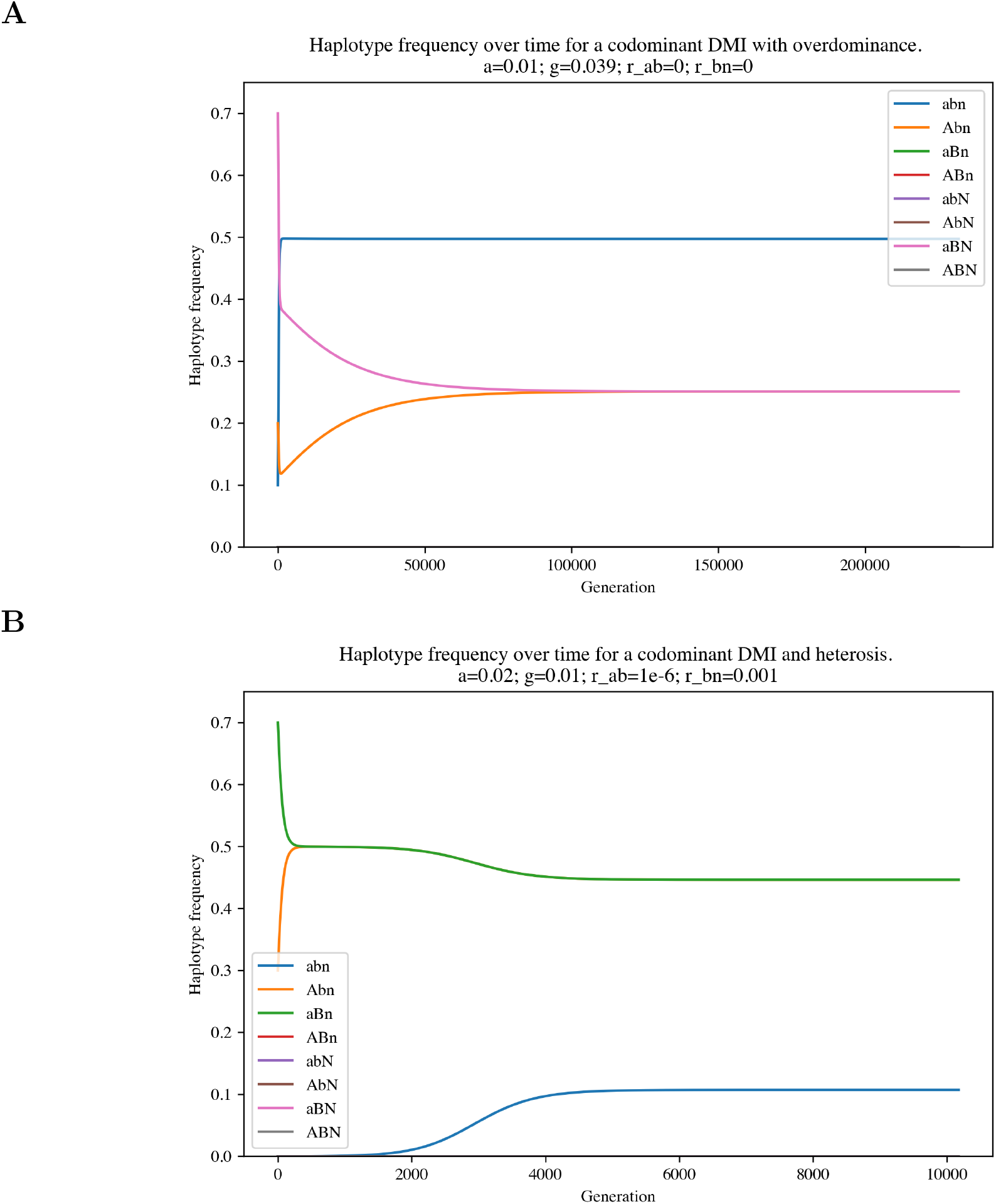
Haplotype trajectory over time until equilibrium is reached on a co-dominant BDMI between co-dominant BDMIs in and, and on neutral . **A** When BDMIs relative to overdominance are strong (*γ* = 0.039 and there is no recombination between *A* and *B*, overdominance with ancestral haplotype *abn* maintains incompatible alleles. 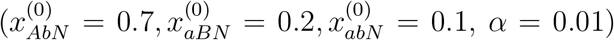. **B** When BDMIs are weak relative to overdominance (*γ* = 0.01) and recombination is low (*r* = 1*e* − 6), parental haplotypes are overrepresented and the ancestral haplotype *abn* is present due to BDMIS. 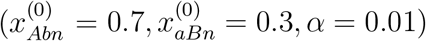

### Linked variation neighbouring a co-dominant BDMI

**Figure S6:**
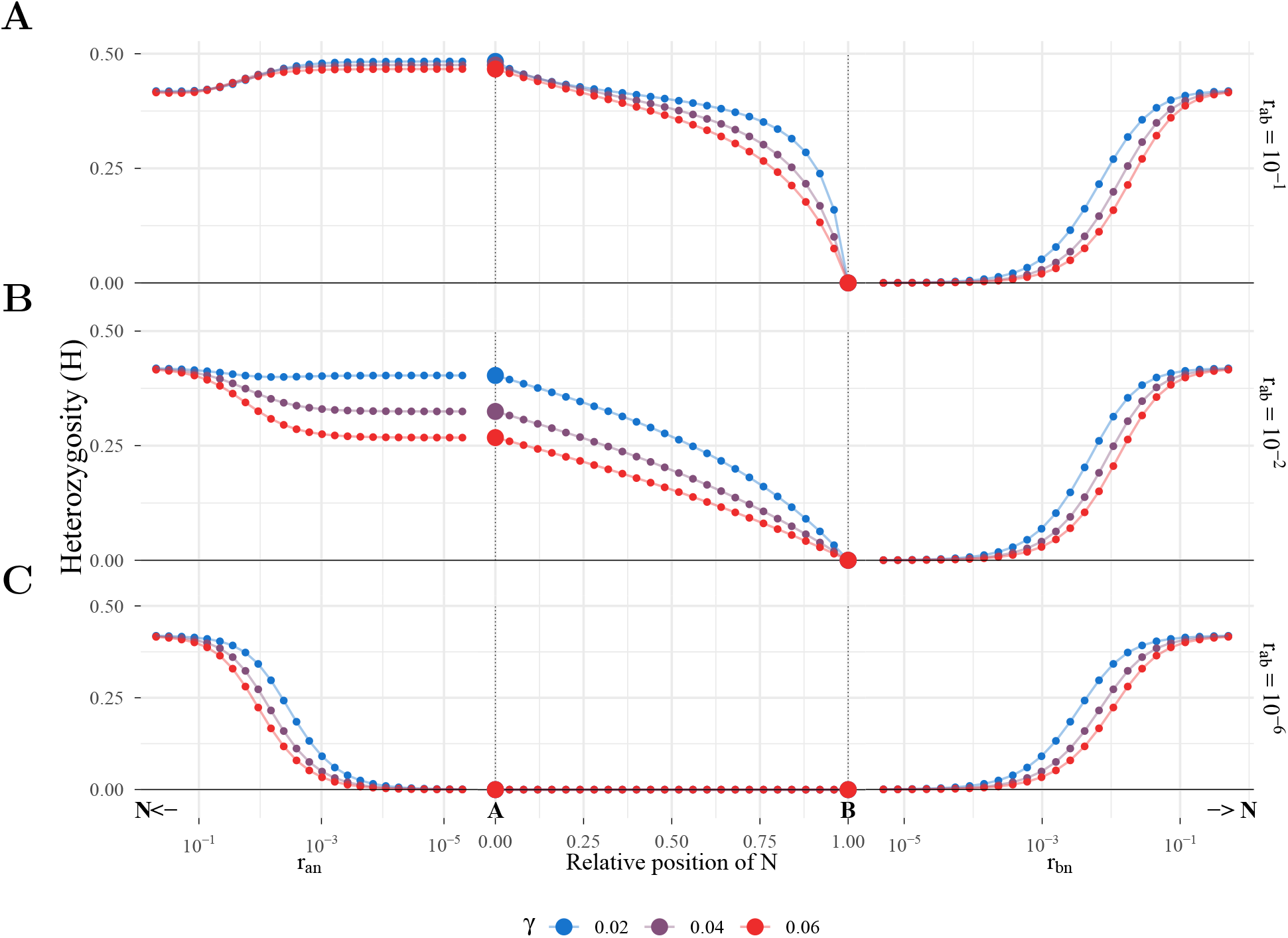
Patterns of heterozygosity linked to a single co-dominant BDMI over different recombination rates (rows).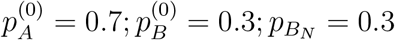

### Effect of initial admixture proportions

**Figure S7:**
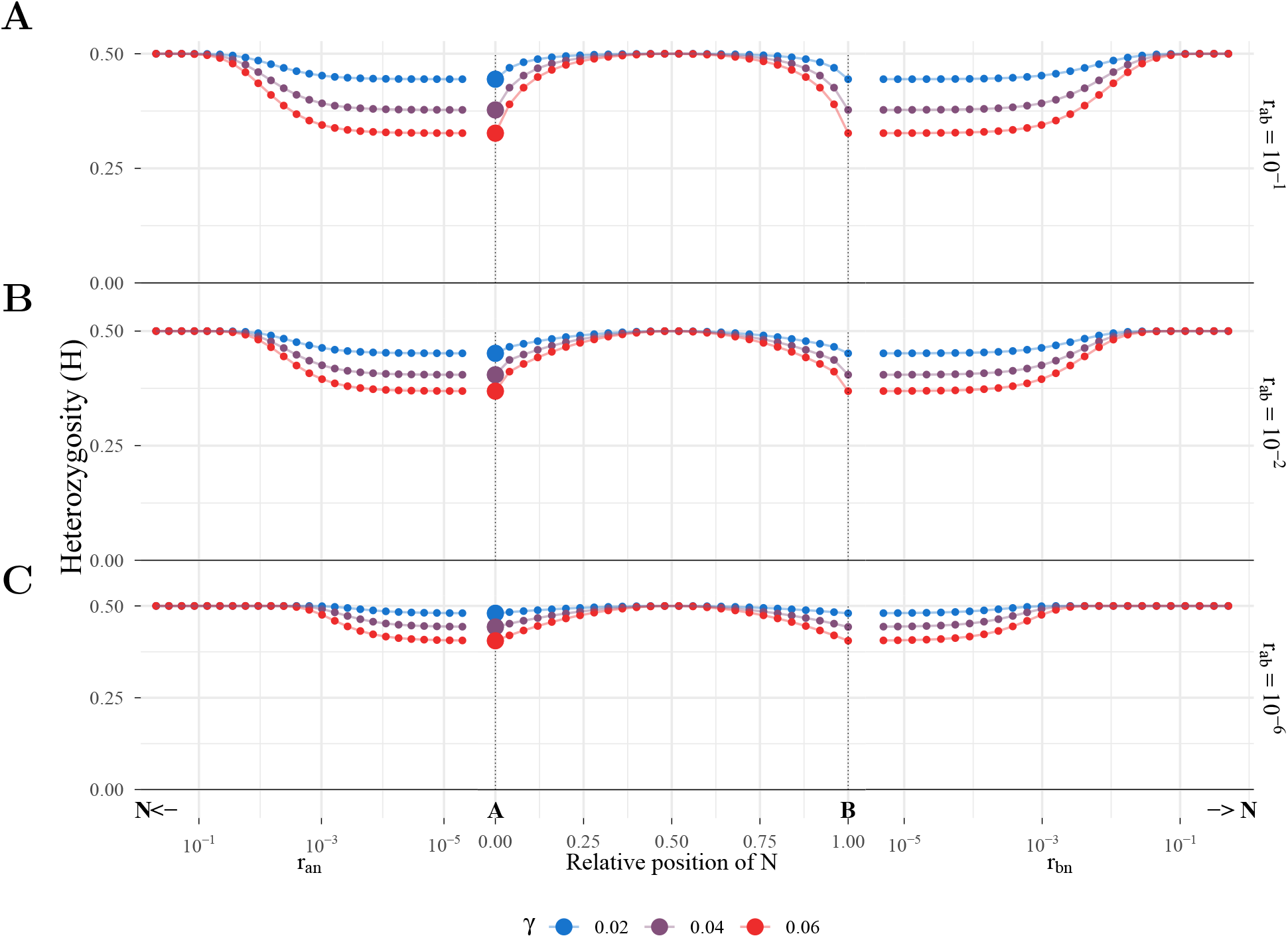
Patterns of heterozygosity at a neutral locus linked to a co-dominant BDMI when incompatible loci *A* and *B* are both overdominant (*α* = *β* = 0.02), with varying recombination *r*_*ab*_ between incompatible loci (high to low, from up to bottom rows). **A** Top row: High recombination (*r*_*ab*_ = 10^−1^) **B** Middle row: Intermediate recombination (*r*_*ab*_ = 10^−2^) **C** Low recombination *r*_*ab*_ = 10^−6^). Initial allele frequencies for all scenarios: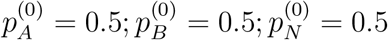;

### Recurrent hitchhiking model fit

To model the reduction in neutral variation caused by the resolution of the BDMI, we leveraged the recurrent hitchhiking model formalised by (Wiehe & Stephan, 1993). Under this model, the expected change in nucleotide diversity is given by:

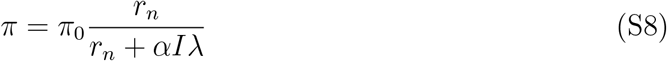

where *π* and *π*0 are the final and initial amount of diversity, *r*_*n*_ is the recombination distance of the neutral locus from the selected site; *α* = 2*Ns* where *N* is the population size and *s*_*rhh*_ the selection coefficient; *λ* is the substitution rate; and *I* is a constant. Since we are interested in a rescaled selection coefficient, we simplify the expression *αIλ* = *s*. Since we are interested in biallelic heterozygosity, *π* = *H*, giving the expression

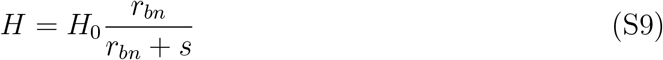

We fit Equation S9 to our numerical solutions for a co-dominant BDMI positioned to the right of the minor allele (Fig. S6, right panels).

**Figure S8:**
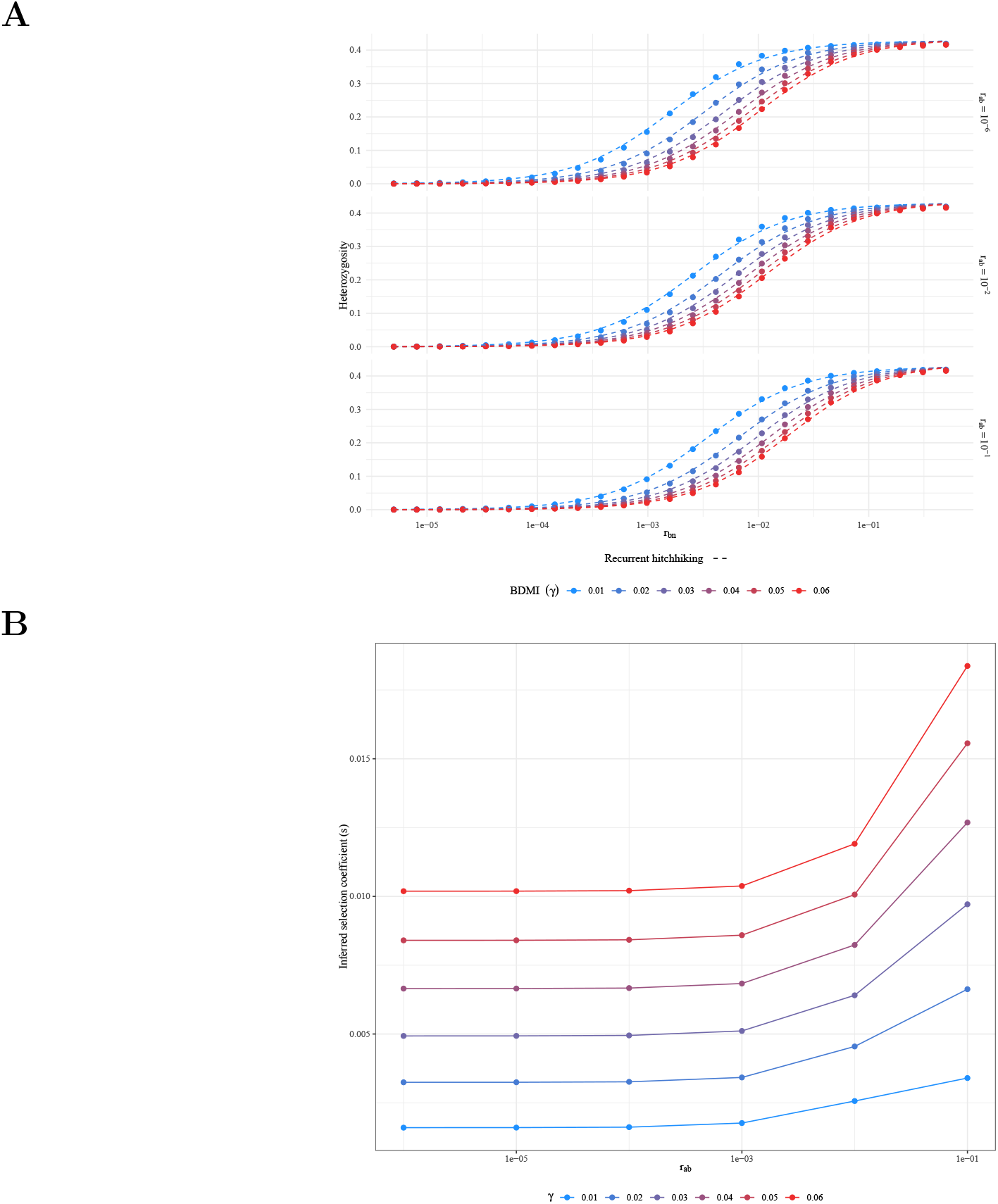
Fit of a BDMI signature of neutral variation to a recurrent hitchhiking model. **A** Observed heterozygosity (points) and fitted model of recurrent hitchhiking (lines) with increasing recombination distance. **B** Inferred selection coefficients of the recurrent hitch-hiking model increase non-linearly with increasing recombination rate between BDMI loci *r*_*ab*_. Co-dominant BDMIs have initial frequencies 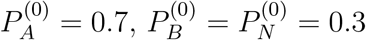

